# Verifying, Challenging, and Discovering New Synapses among Fully EM-Reconstructed Neurons in the Leech Ganglion

**DOI:** 10.1101/427377

**Authors:** Jason E. Pipkin, Eric A. Bushong, Mark H. Ellisman, William B. Kristan

## Abstract

Neural circuits underpin the production of animal behavior, largely based upon the precise pattern of synaptic connectivity among the neurons involved. For large numbers of neurons, determining such “connectomes” by direct physiological means is difficult, as physiological accessibility is ultimately required to verify and characterize the function of synapses. We collected a volume of images spanning an entire ganglion of the juvenile leech nervous system via serial blockface electron microscopy (SBEM). We validated this approach by reconstructing a well-characterized circuit of motor neurons involved in the swimming behavior of the leech by locating the synapses among them. We confirm that there are multiple synaptic contacts between connected pairs of neurons in the leech, and that these synapses are widely distributed across the region of neuropil in which the neurons’ arbors overlap. We verified the anatomical existence of connections that had been described physiologically among longitudinal muscle motor neurons. We also found that some physiological connections were not present anatomically. We then drew upon the SBEM dataset to design additional physiological experiments. We reconstructed an uncharacterized neuron and one of its presynaptic partners identified from the SBEM dataset. We subsequently interrogated this cell pair via intracellular electrophysiology in an adult ganglion and found that the anatomically-discovered synapse was also functional physiologically. Our findings demonstrate the value of combining a connectomics approach with electrophysiology in the leech nervous system.

**Significance Statement:** The function of any nervous system depends on the arrangement of its component neurons into circuits. Determining this precise pattern requires an account of which neurons are linked by synapses, and where. Here we use serial electron microscopy to confirm, challenge, and discover synapses in the neuropil of one ganglion from a juvenile leech. Relying on the homology of the ganglion from animal to animal, we demonstrate that we can identify synapses we knew existed from previous physiological work, and that we can confirm a new anatomically-discovered synapse by subsequently recording from the same neurons in a different animal. Here we show how analyses of anatomical detail and physiologically determined interactions complementarily yield insight into how neural circuits produce behavior.

## Introduction

The behavioral repertoire of a given neural circuit is constrained in part by the connectivity pattern among its constituent neurons. To understand how circuits produce behavior it is therefore necessary to know which neurons make synapses onto which other neurons. Deciphering this connectivity by means of exhaustive electrophysiology is possible in preparations involving relatively few neurons, as in the ~25–30 neuron crustacean stomatogastric ganglion (Marder and Bucher, 2007). As the number of neurons increases however, an imaging-based anatomical approach is required to capture the full connectivity of all neurons within a given volume of tissue (Denk et al., 2012). The resolution necessary to reconstruct neurons and identify synapses among them is provided by serial electron microscopy. For instance, the *C. elegans* hermaphrodite nervous system was reconstructed from a set of overlapping serial electron micrographs from ____ individual animals, resulting in the first whole-organism “connectome” (White et al., 1986). Yet the time-consuming nature of this approach has, until recently, dissuaded attempts to apply serial EM to larger volumes of tissue. In the past decade the development of serial blockface scanning electron microscopy (SBEM; Denk and Horstmann, 2004) and refinement of serial section transmission electron microscopy (ssTEM, e.g. Bock et al., 2011; Ohyama et al., 2015; Kasthuri et al., 2015) has dramatically reduced the image acquisition time for large volumes of neural tissue. The resulting datasets have been used to provide insight into both existing and novel circuits. Among others, these results include discovering new features of a known retinal circuit (Briggman et al., 2011), the circuitry of the tail of male *C. elegans* (Jarell et al., 2012), a new type of retinal bipolar cell (Helmstaedter et al., 2013), the complete visual circuitry of a polychaete worm (Randel et al., 2014), the elucidation of circuits responsible for turning behavior (Ohyama et al., 2015) in larval *Drosophila* as well as olfactory processing in both the larval (Berck et al., 2016, Eichler et al., 2017) and adult *Drosophila* (Tobin et al., 2017, Takemura et al., 2017b), the reconstruction of visual circuits in larval (Larderet et al., 2017) and adult *Drosophila* (Takemura et al., 2013, Takemura et al., 2017a), and the full connectome of the central nervous system of the larval tunicate *Ciona intestinalis* (Ryan et al., 2016).

To link the connectivity information gleaned from SBEM or ssTEM datasets to models of circuit function, the anatomically-predicted synapses must be testable physiologically. In *C. elegans*, the connectome has been essential for guiding cell manipulation, ablation, and functional imaging experiments (Bargmann and Marder, 2013). Similarly, calcium confirmed the existence of synapses identified by EM reconstructions of *Drosophila* circuitry (Ohyama et al., 2015; Takemura et al., 2017a). These applications rely on the ability to identify the same neurons from preparation to preparation – an advantage afforded by many invertebrate systems.

The utility of an anatomically-defined connectivity map is enhanced by the amenability of the preparation to electrophysiological techniques. A connectome specifies which neurons are synaptically connected, but subsequent physiological inquiry is needed to determine whether those connections are inhibitory or excitatory and how strongly a given presynaptic neuron influences its postsynaptic partners. The leech ganglion is particularly advantageous for this purpose as the positioning and size of its neurons render them accessible to sharp electrode intracellular electrophysiology in a way that neurons of *C. elegans* or *Drosophila* are not. In the medicinal leech, *Hirudo verbana*, behaviors are produced by a chain of homologous ganglia each containing approximately 400 neurons. To date, most of the work uncovering the circuitry responsible for given behaviors in the leech has relied on intracellular electrophysiology (e.g. Nicholls and Baylor, 1968; Ort et al., 1974) or optical monitoring of voltage-sensitive dyes (e.g. Briggman et al., 2005). These experiments have resulted in several well-characterized synapses and circuits (e.g. Ort et al., 1974; Stent et al., 1979; Lockery and Kristan, 1990a,b; Kristan et al., 2005), yet many neurons and their connectivity in the leech ganglion remain completely or partly uncharacterized (Wagenaar 2015).

We applied SBEM to leech tissue to study known circuits and discover new synaptic connections. We previously reported on the distribution and pattern of synaptic sites in two SBEM datasets: one small volume of mature leech neuropil, and one entire ganglion taken from the smaller yet behaviorally-mature juvenile leech (Pipkin et al., 2016). Herein, we report on the connectivity uncovered within the juvenile ganglion dataset. To validate the approach, we first analyze the connections of well-characterized motor neurons that innervate the longitudinal muscles and participate in the swimming behavior. Second, we use the dataset to identify a previously uncharacterized synaptic relationship and subsequently verify it physiologically. Our results demonstrate the utility and potential of EM-based circuit reconstruction in the medicinal leech by linking anatomy and electrophysiology at the level of individual cell pairs.

## Materials and Methods

### Animals

We used both adult and juvenile medicinal leeches (*Hirudo verbana*). Adult leeches were obtained from Niagara Leeches (Niagara Falls, NY) and housed in aquaria on 12 h daily light/dark cycle at 15–16°C. Juvenile leeches were obtained by harvesting cocoons produced by a breeding colony of adult leeches maintained in our laboratory. Leeches were allowed to mature within the cocoons at room temperature (RT) and collected once they had emerged. We then waited two weeks to ensure full development prior to dissection. We confirmed that the juveniles lacked any embryonic features using established staging criteria (Reynolds et al., 1998). For the juvenile sample, we stained and embedded several ganglia but eventually imaged only ganglion 11. The methodological description of this sample’s preparation (below) and results of some analyses have been published previously (Pipkin et al., 2016).

### Sample preparation for electron microscopy

We anesthetized the juvenile leech in ice-cold leech saline (4°C) containing 115mM NaCl, 4 mM KCl, 1.8mM CaCl_2_, 2mM MgCl_2_, 10mM HEPES buffer (Nicholls and Purves, 1970). Midbody ganglia were then dissected from the nerve cord and pinned to the bottom of a Sylgard-coated dish. The ganglia were then fixed for two hours at RT in 2% PFA, 2.5% glutaraldehyde, and 0.1M phosphate buffer. After fixation the ganglia were rinsed in 0.1M phosphate buffer and incubated in 2% OsO_4_ / 1.5% potassium ferrocyanide. For this step, the samples were microwaved in a scientific microwave (Pelco 3440 MAX) three times at 800 W with a duty cycle of 40 seconds on and 40 seconds off at a measured temperature of 35°C and subsequently left to sit at RT for thirty minutes. Samples were then washed in ddH_2_O and microwaved three times at 800 W with a duty cycle of 2 minutes on and 2 minutes off at 30°C. We found that this and subsequent brief microwave incubations facilitated staining penetration to the center of our samples and was necessary to gain sufficient image contrast. Samples were then incubated in 1% thiocarbohydrazide (Electron Microscopy Sciences) and microwaved three times at 800 W with a 40 seconds on and 40 seconds off duty cycle at 30°C and subsequently left to incubate for 15 minutes RT. The samples were then washed again with the same microwave incubation as described earlier. Next, the samples were incubated in 2% aqueous OsO_4_ and microwaved three times at 800 W with a 40 seconds on and 40 seconds off duty cycle at 30°C and then incubated at RT for one hour. After washing, the samples were then left in 1% uranyl acetate overnight at 4°C. The next day, samples were incubated in a lead aspartate solution prepared by dissolving 0.066 gm of lead nitrate into 10 ml of 0.03M aspartic acid with the pH subsequently adjusted to 5.5 using 1N KOH. This incubation took place in a 60°C oven for 30 minutes. The samples were then washed and dehydrated through a series of ethanol solutions (50%, 70%, 90%, 100%, 100%, 10 minutes each) at RT and incubated in acetone. Following this, samples were infiltrated with epoxy resin by first incubating them for two hours at RT in a solution of 50% acetone and 50% Durcupan and then overnight in 100% Durcupan. The next day, samples were transferred to a freshly prepared 100% Durcupan solution and incubated at RT for 2 hours. Samples were then incubated within a 60°C oven for three days. Durcupan Araldite resin was made by mixing 11.4 g of component A, 10 g of component B, 0.3 g of component C, and 0.1 g of component D.

### Imaging

The resin-embedded ganglia were preserved within epoxy blocks trimmed until tissue was barely exposed. For the juvenile sample, the edges of the block were trimmed until very near to the external capsule of the ganglion to reduce charging in the outer image tiles that contained both tissue and empty plastic. Blocks were mounted onto aluminum pins to which they were adhered with conductive silver paint. The pin and block were then sputter coated with a thin layer of gold and palladium to further enhance conductivity.

We imaged ganglion 11 from a juvenile animal with a Zeiss MERLIN SEM equipped with a Gatan 3VIEW SBEM system. We collected montages of 8000x8000 raster tiles at 5.7 nm pixel size. We oriented the sample so that it was imaged from the dorsal surface to the ventral surface with sectioning occurring perpendicular to the dorsal-ventral axis. Montage size thus varied from 1x1 to 5x5 tiles depending on the area of tissue that was exposed to the surface of the block. We sectioned the block 2203 times at 50, 100, or 150 nm thicknesses for a total z-distance of 138 μm. The 100 nm and 150 nm sections were taken in regions containing only cell bodies (at the top and bottom of the overall volume) as there are very few fine neuronal processes to trace here and thus imaging time could be reduced. Similarly, we varied dwell time throughout acquisition along a range of 0.8-μs to 1.5-μs with higher dwell times used in neuropil-containing sections. During the juvenile ganglion acquisition, an unexpected and gradual reduction of contrast occurred due to the premature degradation of the filament in the electron gun. As imaging proceeded from the dorsal surface towards the ventral, we therefore focused most of our analysis and reconstruction on cells whose arbors tended to fall within the dorsal half of the ganglion. Where processes from these cells entered the low-contrast region of the volume, we were likely to have missed some fine branches and any associated synapses in this area.

### Reconstruction and Annotation

In the juvenile ganglion volume, montages and sections were aligned in the TrakEM2 (fiji.sc/TrakEM2, RRID: SCR_008954; Cardona et al., 2012). Subsequent tracing and annotation was also performed in TrakEM2. In this volume we largely reconstructed arbors via skeletonization rather than full segmentation via membrane tracing. Locations of synaptic inputs and outputs were denoted by placing ball objects as markers on the skeletons.

All tracing, segmentation, and analysis was performed by JP. To reduce errors, the arbors of the motor neurons discussed in Table 1 and Figures 1 and 2 were reviewed at least twice. As has been previously reported (Ohyama et al., 2015), false negatives (missing branches) were far more likely errors than false positives (adding the wrong branch).

**Figure 1.**
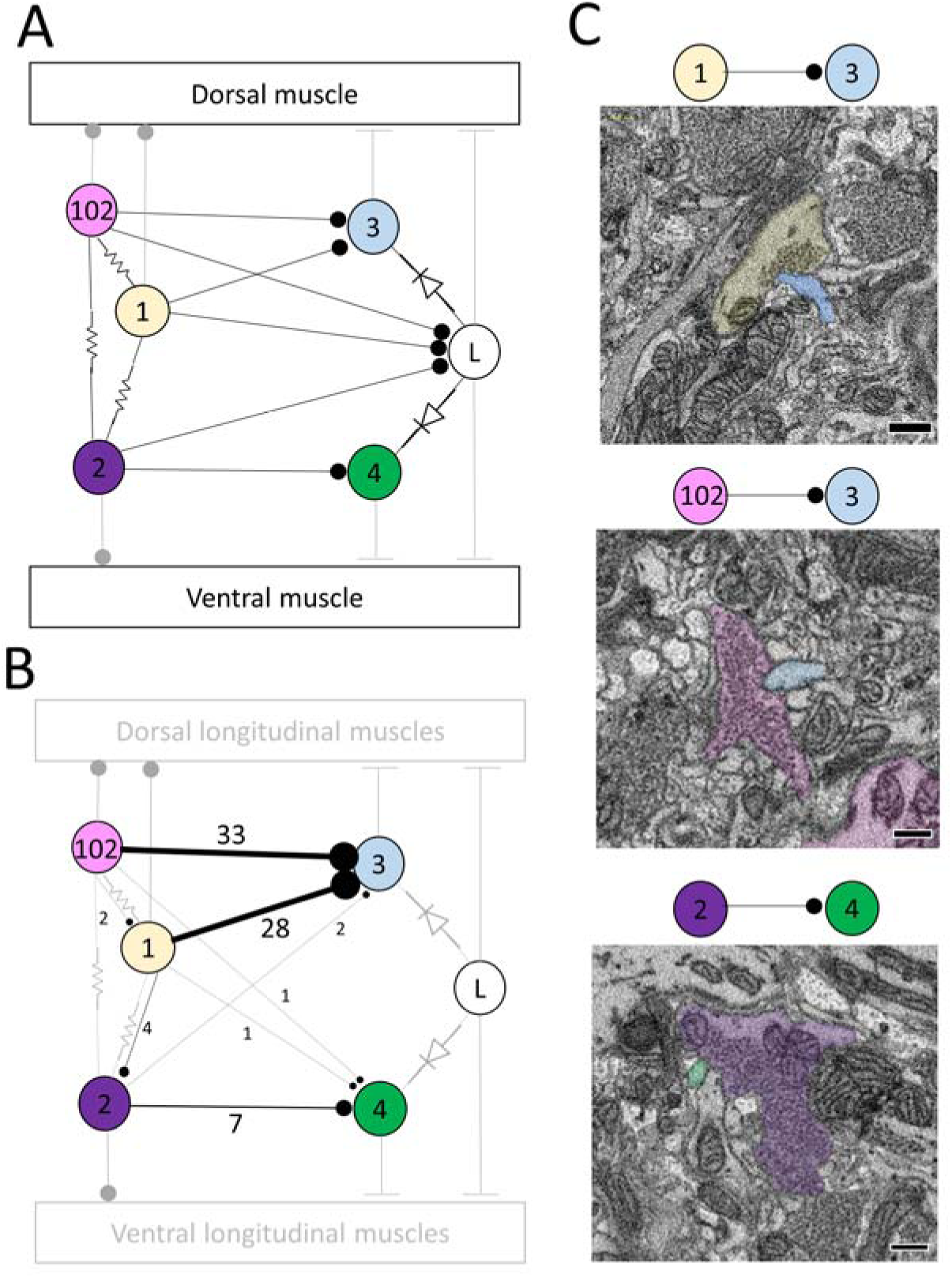
Most, but not all, of the predicted physiological connections were found anatomically after reconstructing the arbors of six pairs of dorsal motor neurons. **(A)** Predicted circuitry based on dual electrophysiological recordings, adapted from Ort et al. (1974). Lines ending in circles represent inhibitory connections; lines ending in a T-junction indicate excitatory connections; resistors indicate non-rectifying gap junctions; diodes represent rectifying gap junctions. **(B)** Updated circuitry based on what was directly observed after anatomical reconstruction. Electrical connections are grayed out as these are not directly observable with SBEM. All predicted connections were found except those onto the L cell. A few unexpected synapses were found (e.g. from cell 1 to cell 102); these typically involved far fewer overall synapses (Table 1). The total number of synaptic contacts made by both the right and left pairs of neurons are shown next to each line (see also Table 1). **(C)** Examples of synapses between the right DI-1 and the right DE-3 (upper panel), the right DI-102 and the right DE-3 (middle panel), and the left VI-2 and the right VE-4 (lower panel). In these examples, the cells are fully segmented to display the relative scale of the participating processes; the remainder of their arbors were traced via skeletonization. Scale bars 300nm.

**Figure 2.**
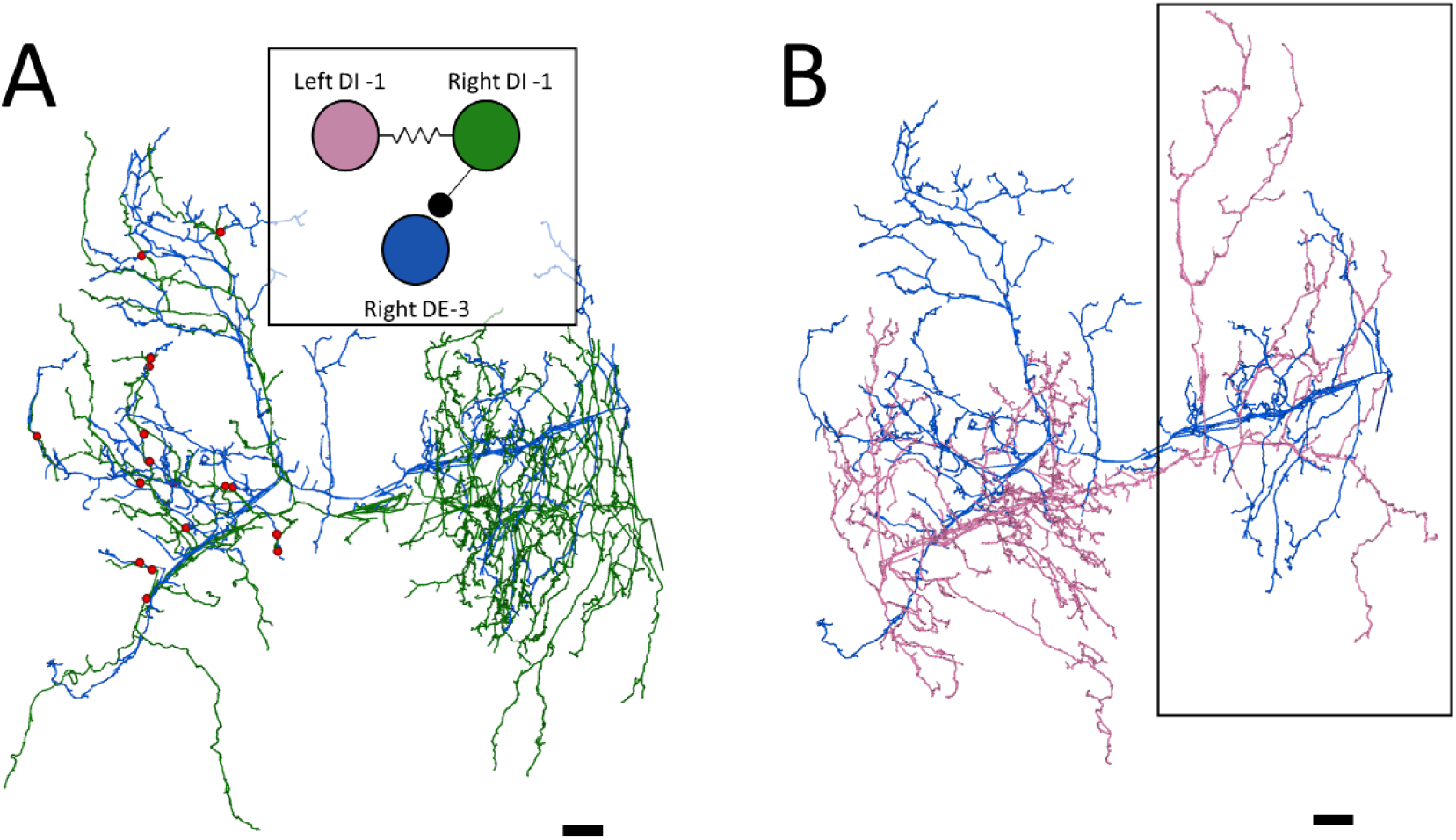
The right DE-3 receives numerous widely-distributed synaptic inputs from the right DI-1 and none from the left DI-1. **(A)** The right DE-3 (blue skeleton) receives synaptic input from the right DI-1 (green skeleton) at 18 sites (red dots) widely distributed throughout the contralateral half of its arbor. Inset displays the previously-known connectivity among these three cells. **(B)** The left DI-1 arbor (pink skeleton) overlaps with the right DE-3 arbor. Even where the left DI-1 forms presynaptic boutons and the right DE-3 receives synaptic inputs, no synapses are found (region within black box). 10μm scale bars. Arbors are presented as viewed from above, with anterior to the top. Cell bodies are omitted for clarity, as their position above the arbors would partially obscure them.

**Table 1.**
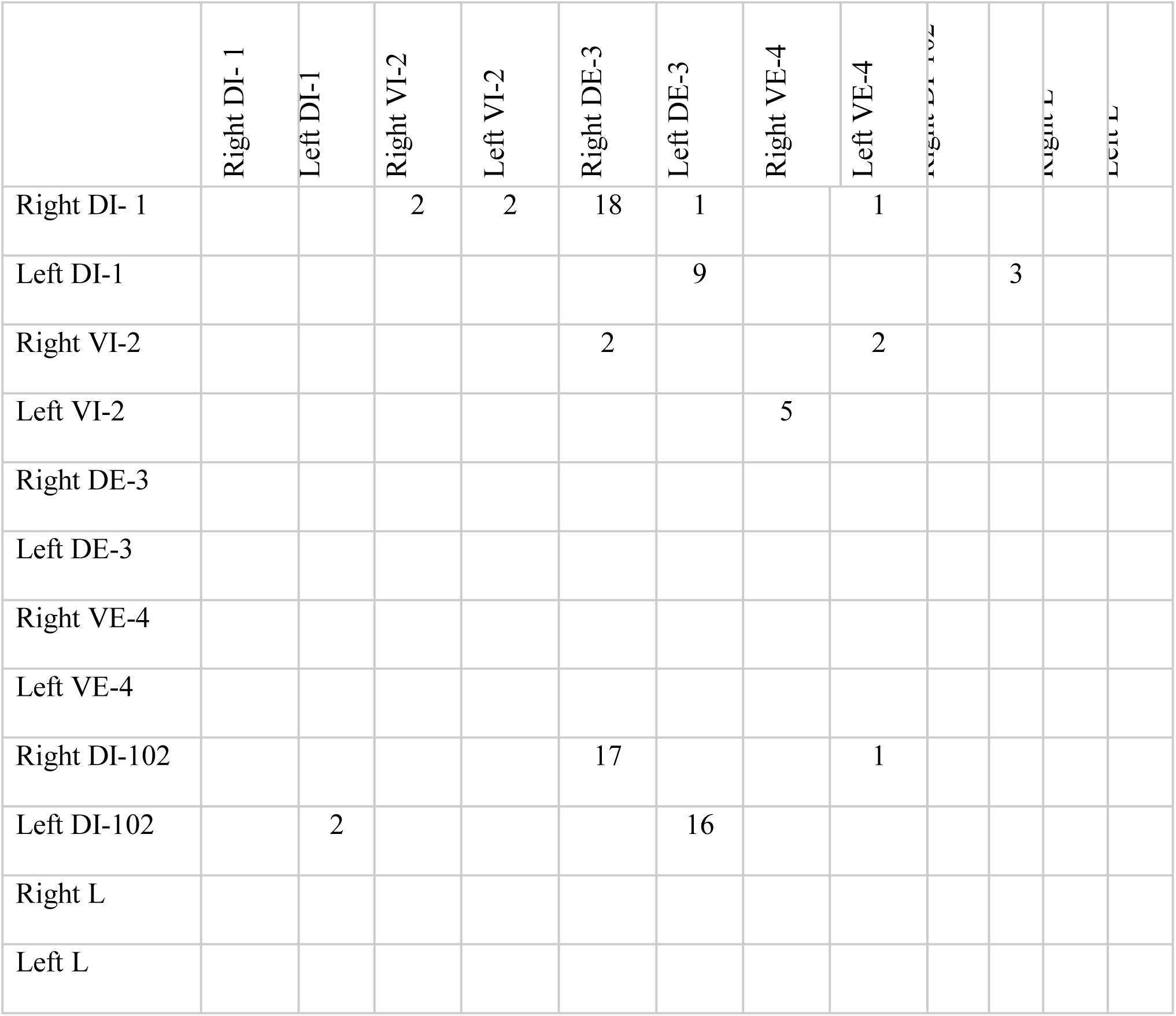
Number of chemical synaptic contacts found among six pairs of motor neurons. Presynaptic cells are listed in the first column and postsynaptic cells are listed in the first row. All expected connections were found, with the exception of direct connections from DI-1, DI-102, or VI-2 onto the L cells. Some unexpected synapses were also found but were typically low in number compared to expected synapses (e.g. right DI-1 onto left DE-3).

### Electrophysiology

Adult leeches were anesthetized in ice-cold saline, dissected, and chains of four midbody ganglia were removed and pinned in Sylgard-coated dish. The ventral sheath of the second ganglion and dorsal sheath of the third ganglion were removed to expose cell bodies for penetration with 1.0 mm (OD) X 0.75 mm (ID) glass microelectrodes with an omega dot pulled to a resistance of ~20 MΩ. Microelectrodes were filled with 20 mM KCl and 1 M potassium acetate. To verify that the S cell was impaled, and its intracellular spikes were matched 1:1 with the largest extracellularly-recorded spikes in the connective between the third and fourth ganglia. To verify cell 116’s identity, we loaded electrode tips with either Alexa Fluor 488 or Alexa Fluor 594 (Thermo Fisher Scientific) and filled the electrode shanks with 3M potassium acetate. Dye was then injected with alternating depolarizing and hyperpolarizing current pulses (2nA for 300 ms, -2nA for 50ms, 10% duty cycle for 30 minutes) and the shape of the arbor compared to the reconstructed arbor from the juvenile ganglion SBEM dataset.

Intracellular current injection and measurement of membrane potential were mediated by an Axoclamp-2B amplifier (Axon Instruments, Inc.) operated in bridge mode. Extracellular recordings were amplified by a Model 1700 A-M Systems differential amplifier. Electrical signals were digitized, recorded, and analyzed with WinWCP (Strathclyde Electrophysiology Software). Further analysis was performed with Microsoft Excel (Microsoft).

### Experimental Design and Statistical Analysis

This bulk of this work represents analyses of a single volume of electron micrographs containing one juvenile leech ganglion. We therefore do not make any statistical comparisons – we present our work as a set of observations which can then be compared to prior work and suggest future experiments. Similarly for our electrophysiology experiment, we do not make any statistical comparisons.

## Results

### Neuron and synapse identification

The somata of leech neurons are arrayed along an outer rind of each midbody ganglion. To identify a soma in our EM volume, we first compared its size and location with the known map (Ort et al., 1974; Muller et al., 1981). Soma position can vary slightly from ganglion to ganglion, but the basic shape of the neuron’s arbor can distinguish it from its neighbors (Fan et al., 2005). Our identifications were based on a combination of soma size, position, and arbor morphology. By convention, neurons are named according to their corresponding letter or number identifier in the accepted map (Ort et al., 1974; Muller et al., 1981). In the case of motor neurons, these cell number identifiers are preceded by two letters indicating which motor group the cell innervates and whether its outputs are excitatory or inhibitory. For example, cell DI-1 is an inhibitor of the dorsal longitudinal muscles while cell VE-4 is an excitor of the ventral longitudinal muscles. Most neurons in the leech ganglion are paired, having both a right and a left homologue. An exception (the “S-cell”) is considered below.

We identified synapses by the criteria discussed in recent work (Pipkin et al., 2016). Briefly, leech presynaptic varicosities lack densely-staining T-bars characteristic of neuropil in *Drosophila* and some other invertebrate preparations. Instead, presynaptic sites are labeled by small presynaptic tufts of electron-dense material and faint postsynaptic densities that are indistinguishable at the resolution afforded by SBEM (Purves and McMahan, 1972; Muller and McMahan, 1976; Muller and Carbonetto, 1979). Our requirements for synapse identification were twofold: (1) a concentration of small presynaptic vesicles in the presynaptic neuron, some of which lie near to the membrane apposition of presynaptic and postsynaptic neurons; and (2) that the apposition of presynaptic and postsynaptic membranes persists over three or more consecutive imaging sections. Our criteria are more liberal than those afforded by higher-resolution TEM. As a result, while they capture the majority of real synapses we cannot exclude the possibility that we have mis-identified some nonsynaptic appositions as synapses.

### Testing physiologically-characterized circuits anatomically

The synaptic connections among neurons that generate behaviors in the leech are made in the neuropil of each ganglion. Within our juvenile ganglion volume, we explored the connections of a subset of motor neurons known to participate in the swimming behavior (Ort et al., 1974). Specifically, we searched the neuropil for synapses among the bilateral pairs of neurons DI-1, VI-2, DI-102, DE-3, and VE-4, which innervate dorsal and ventral longitudinal muscles and are responsible in part for the undulation of the leech’s body during swimming. In addition, we also searched for connections made by these cells with the pair of L motor neurons, which are excited during the shortening reflex but are inhibited throughout swimming.

The physiologically-determined circuit among these cells is depicted in Figure 1A. (adapted from Ort et al., 1974). In this diagram, non-rectifying electrical synapses are represented by resistors and rectifying electrical synapses are represented by diodes. As the resolution of SBEM precludes the direct observation of gap junctions, we turned our attention first to chemical synapses (Figure 1A). We first sought to locate and quantify the number of known inhibitory synapses made within the neuropil in this circuit. To do so, we manually traced skeleton arbors of all the neurons involved, noting where each neuron made a synapse onto the other neurons (Figure 1B,C), using the criteria established in our previous study (Pipkin et al., 2016). The number of synapses formed in this network are summarized in the connectivity matrix shown in Table 1. We found numerous synaptic contacts consistent with the previously-described direct inhibition of DE-3 by the ipsilateral DI-1 and DI-102 and the direct inhibition of VE-4 by the ipsilateral VI-2. We did not find any chemical synapses from DI-1, DI-2, or DI-102 onto either L cell (Figure 1B), suggesting that the observed physiological inhibition occurs via an indirect pathway, potentially via the electrical connections.

As suspected from electrophysiological recordings (Ort et al., 1974), we observed that each DE-3 received direct inhibitory input from the ipsilateral DI-1. We previously observed that each DI-1 forms presynaptic boutons only in the contralateral portion of their arbors (Pipkin et al., 2016). In Figure 2A, the right DI-1 (green) is presynaptic to the right DE-3 via 18 synapses (red dots). Within the contralateral arborization of DE-3, these 18 synapses were widely distributed, contradicting previous predictions that inputs from DI-1 might be concentrated onto a single branch (Lytton & Kristan, 1989). We found a similar pattern among the inputs from the DI-102s onto the DE-3s (data not shown). Notably, the right DE-3 received no input from the left DI-1, despite overlap of the vesicle-containing portion of the left DI-1’s arbor with the ipsilateral arborization of the right DE-3 (box, Figure 2B). With the exception of a single synaptic contact, this was also true for the right DI-1 and left DE-3 and for both DI-102s and DE-3s (Table 1). Similar to the dorsal muscle inhibitory motor neurons (DI-1 and DI-102), the ventral inhibitor (VI-2) neurons form presynaptic boutons in only the contralateral portion of their arbor. Consistent with the fact that the pair of ventral excitatory motor neurons (VE-4) arborize exclusively in the in the ipsilateral half of the neuropil each VE-4 received direct inhibition only from the contralateral VI-2 (Table 1), a finding that agrees with the electrophysiological characterization of this connection (Ort et al., 1974).

### Electrical connections

It is impossible to directly observe the fine structures characteristic of gap junction membrane appositions when constrained by the resolution limits of SBEM (Brightman and Reese, 1969). Nonetheless, several of the cells we traced formed electrical connections with each other on the basis of prior electrophysiological evidence (Ort et al., 1974). We therefore took note when the membranes of two cells known to be electrically-coupled came into extended contact over many sections. On the basis of this criterion, we observed several suggestive contacts. In some cases, the contact is extensive in area and seen at many separate sites. For example, we traced the S cell, a unique excitatory interneuron involved in the shortening reflex (Laverack 1969; Frank et al., 1975; Magni and Pellegrino, 1978; Crisp and Muller, 2006) and known for its large fast-conducting axon that it extends both anteriorly and posteriorly in Faivre’s nerve. Halfway between each ganglion, this axon forms an electrical synapse with the S cell of the adjacent ganglion such that spikes generated in one S cell are propagated throughout the entire nerve cord (Muller and Carbonetto, 1979). Additionally, the S cell is known for making strong electrical connections with two “coupling interneurons” that act in part as relays for sensory inputs (Muller and Scott, 1981). In Figure 3A, we show a confluence of processes belonging to the S cell (blue) and one of each coupling interneuron (green and pink). In this particular junction, each cell’s membrane is closely apposed to and conforms to each other’s and this interaction persists over several sections. We also searched for contacts among other known coupled cells. For example, Figure 3B depicts the close apposition of the left DI-102 (red) and left DI-1 (yellow). Both these cells are known to be physiologically coupled (Figure 1A). Here two of their secondary branches come into close contact as they travel adjacent to each other; notice again that both cells’ membranes are closely apposed and conformed to each other. Not all possible junction sites involved symmetrically sized processes. In one case, a thin process belonging to the left DE-3 (orange) burrows into the primary process of the right DE-3 (purple) (Figure 3C). Again, both these cells are known to be coupled (almost all pairs of dorsal motor neurons are electrically coupled [Ort et al., 1974; Fan et al., 2005]). In every instance involving known electrically coupled cells, we observed sites of membrane contact that could harbor gap junctions. For instance, we found 24 and 26 contacts between the S cell and each coupling interneuron, 5 between the left DI-1 and left DI-102, and 10 between both DE-3s. Like chemical synapses (Figure 2A,B), these contact sites were distributed throughout cells’ arbors. Because we traced arbors chiefly via skeletonization, we cannot say whether the cumulative amount of membrane apposition predicts electrical connections. However we can report that we did not observe similarly prolonged, conformed appositions among cells not known to be coupled.

**Figure 3.**
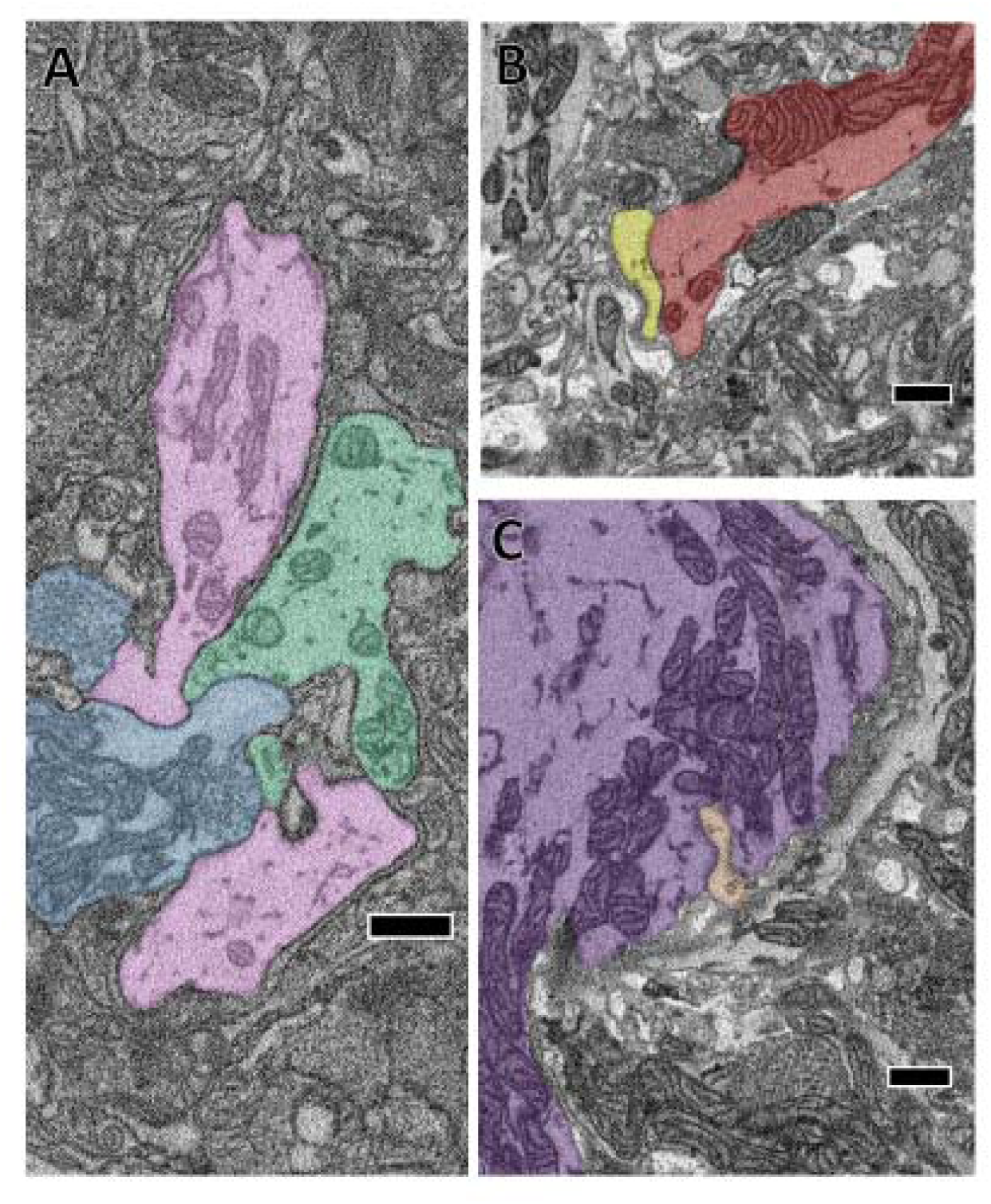
The close apposition of cell pairs known to be electrically coupled could harbor gap junctions. Though the arbors shown were all traced by skeletonization, we fully segmented them in each particular section shown here to highlight their membrane appositions. **(A)** The confluence of the S cell (blue) and both coupling interneurons (pink and green). **(B)** Close apposition between two processes of the left DI-102 (red) and left DI-1 (yellow). **(C)** A small branch of the left DE-3 (orange) invades the main branch of the right DE-3 (purple). Scale bars 500nm.

### Predicting a physiological connection from an anatomical connection

We next sought to test whether an anatomical synapse predicts a physiological connection. For this experiment, we turned to cell 116. Each cell 116 is inhibitory and resides in the dorsal aspect of the anterolateral packet (E.P. Frady and K. L. Todd, personal communication). In tracing arbors of the pair of cells 116, we noticed that each received synaptic input from the S cell. The S cell (blue skeleton, Figure 4A) made 6 synapses onto the right 116 (orange skeleton, Figure 4A) and 7 synapses onto the left 116 (green skeleton, Figure 4A), distributed throughout the extent of the S cell arbor (pink dots, Figure 4A). In one case, both cells 116 were postsynaptic to the same S cell bouton.

**Figure 4.**
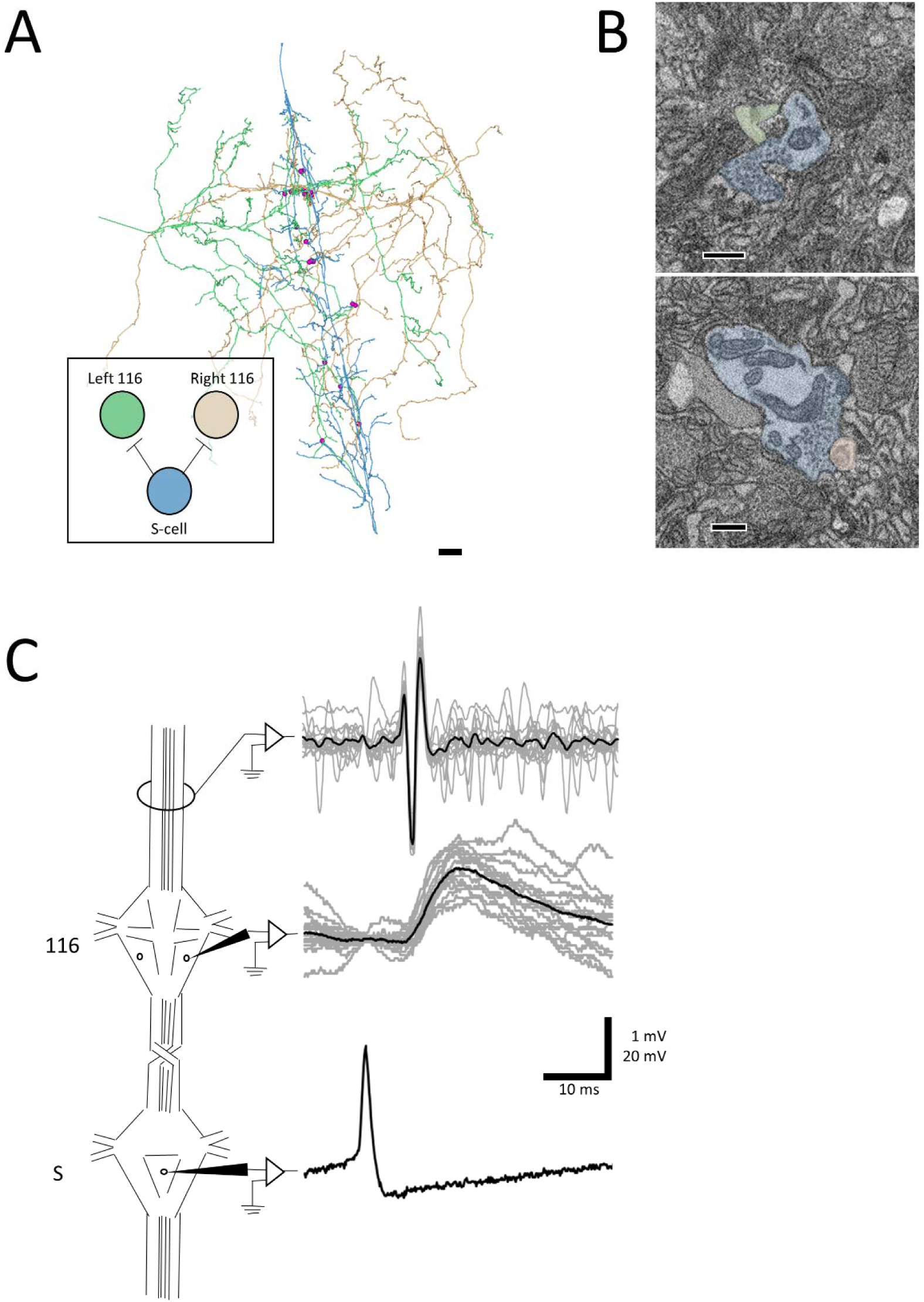
A synapse discovered anatomically makes an electrophysiological connection. **(A)** Skeleton arbors of the presynaptic S cell (blue) and postsynaptic cells 116 (green and orange) with pink dots representing sites of synaptic contact. 10μm scale bar. Arbors are presented as viewed from above, with anterior to the top. Cell bodies are omitted for clarity, as their position above the arbors would partially obscure them. Inset displays the connections between the S cell and cells 116 that we tested physiologically. **(B)** Examples of synapses from S onto the left 116 (top) and right 116 (bottom). 300nm scale bars. Cells are fully segmented in these example sections to display the relative scale of the individual processes; the remainder of the arbors were reconstructed via skeletonization as shown in (A). **(C)** Example recordings from one adult nervous system preparation of the S-116 connection. Spikes were induced in the S cell in one ganglion (bottom trace) whereupon they traveled across the S cell network down the nerve cord, eliciting a reliable depolarization in cell 116 (middle trace). The S cell spike was visible in an extracellular recording of the connective nerves posterior to the ganglion containing the recorded 116, indicating that the spike successfully passed through (top trace). A single spike in the S cell is presented for clarity in the bottom trace while the middle and top represent recordings following 15 separate S-cell spikes from the same preparation (grey) and their average (black).

We next tested to see if inducing action potentials in the S cell network would reliably lead to excitatory potentials in cell 116. Because the S cell in one ganglion is strongly coupled to the S cells in the next ganglion anterior or posterior to it, we were able to circumvent the practical difficulty of simultaneously recording intracellularly from one cell on the ventral surface and another cell on the dorsal surface. Instead, we impaled the S cell in the ganglion adjacent to the one in which we recorded cell 116 (Figure 4C). To confirm that the spike traversed through the network, we recorded the connective nerves posterior to the cell 116 ganglion with an extracellular electrode (Figure 4C). We observed that each S cell spike reliably preceded a 1–2 mV EPSP in cell 116. The cell 116 response to 15 S cell spikes (overlaid, grey traces in middle panel) is presented in Figure 4C along with their average (black trace in middle panel). The 4–5ms latency between spike and EPSP is consistent with known conduction velocity of the S cell spike through Faivre’s nerve (Frank et al., 1975).

## Discussion

Our results validate a connectomics approach for circuit discovery in the leech ganglion. We show that reconstruction of selected cells can be used to confirm the existence of previously known connections among motor neurons (Figure 1, Table 1). Previous work showed that the ipsilateral DI-1 and DI-102 monosynaptically inhibit DE-3, while the contralateral VI-2 inhibits VE-4 (Ort et al., 1974; Granzow et al., 1985). At the resolution of light microscopy, others have observed considerable overlap between the processes of these cells and have noted possible sites of apposition of postsynaptic processes with presynaptic varicosities (Granzow et al. 1985, Fan et al., 2005). At the EM level, Granzow et al. (1985) attempted to demonstrate the connection between DI-1 and DE-3 by differentially staining the two cells with intracellular fills (Imposil in DI-1, horseradish peroxidase in DE-3) and taking thin sections of the contralateral half of the neuropil. However, due to suspected disruption of vesicle structure wrought by Imposil they found presynaptic vesicles near only one of many sites of abutment between the two cells (Granzow et al., 1985). By analyzing a complete SBEM volume of an entire ganglion, our report is the first to provide direct EM anatomical confirmation of these synapses among motor neurons.

For each of these known connections (DI-1 -> DE-3, DI-102 -> DE-3, VI-2 -> VE-4) we found more than one synapse from the presynaptic cell onto the postsynaptic cell. The number of such contacts ranged from 2 (from the right VI-2 onto the left VE-4) to 18 (from the right DI-1 onto the right DE-3) (Table 1). It is unclear in what ways this variability is physiologically meaningful, as we cannot infer the synaptic strength of a given synapse in a SBEM volume. While it is tempting to speculate whether a connection with more contacts is stronger than one with fewer, the highly electrically-coupled motor circuit we reconstructed is not ideal for addressing this question. Other subcircuits in the leech ganglion are more attractive. For example, the connections among the sensory P cells and local bend interneurons are known to vary in strength physiologically (Lockery and Kristan, 1990a). Unfortunately, the cells involved primarily form their arbors in the ventral aspect of the neuropil, where the deteriorated quality of our dataset precluded accurate reconstruction (see Materials and Methods). Ongoing work carrying this project forward in the adult leech ganglion by the Wagenaar and Ellisman groups may be able to more fully explore the relationship between contact number and physiological synapse strength.

The range of contact number we observe falls below that measured by light microscopic analysis of overlap between adult sensory and motor neurons (13–41 in DeReimer and Macagno, 1981). This difference could be due to the maturity of the tissue, the specific cell pairs studied, or methodological differences (processes may overlap at the light level but do not touch at the EM level). The range of synapse number per connection that we find (1–18) is somewhat comparable to what has been found in other systems in which entire arbors have been reconstructed from serial EM images (*C. elegans*: 1–19 in the hermaphrodite [J.G. White et al., 1986], 1–61 in the male tail [Jarrell et al., 2012]; *Platynereis dumerilii*: 1–45 including neurons and muscles of visuomotor circuitry [Randel et al., 2014]; *Drosophila melanogaster:* 1–199 in the visual circuitry [Takemura et al., 2013], 1–23 from a selectively reconstructed motor circuit in the larva [Ohyama et al., 2015]).

We observed some unexpected sites of potential synaptic contact among the motor neurons we traced (for example, the right DI-1 makes a single synapse onto the left VE-4, Table 1). Notably, these cases involve far fewer overall contacts (1–3). Ohyama et al. (2015) also examined circuitry in which multiple types of the same cell in *Drosophila* larvae (Basins 1–4) made inputs onto various postsynaptic cells. In their data, they report instances where these postsynaptic cells predominantly receive input from one of these Basin cell types while still receiving scattered input from the others (for example, the cell they label A12q a1l receives 15/14 synaptic inputs from the right/left Basin 2s and 0/1 from the right/left Basin 1). There are a number of possible explanations for our finding of unexpected connections: (1) these synapses may be real but so relatively few in number as to be physiologically undetectable and unimportant; (2) these synapses may be present only in juvenile tissue that is still undergoing synaptic refinement; (3) these synapses could be mistakenly identified or otherwise be the result of a tracing error that we cannot detect after reviewing them; (4) some of these synapses might actually be gap junctions occurring at a location that makes them appear to be chemical synapses (e.g. the connections between left DI-1 and left DI-102, two inhibitory neurons known to be electrically coupled [Fan et al. 2005]).

We found that synapses between two cells widely spanned the region of overlap between the vesicle-containing portion of the presynaptic cell’s arbor and the postsynaptic cell’s arbor (Figure 2). Earlier reports had suggested that the synapses made by DI-1 and DI-102 might be concentrated onto separate single branches of the DE-3 arbor (Lytton and Kristan, 1989). We find no evidence for such selectivity in our juvenile ganglion volume, though we cannot rule out that synapse strength might vary depending on where a synapse occurs or that branch-selectivity is a process that is not yet complete in juvenile tissue.

We almost exclusively found synapses from the DI-1 and DI-102 cells onto the ipsilateral DE-3 even though the vesicle-containing portion of the DI-1 or DI-102 arbor overlaps with postsynaptic regions of both the ipsilateral and contralateral DE-3. This lateral selectivity suggests that there may be some chemical basis by which synapse formation is restricted to the ipsilateral cell pair. This result also underscores the strengths of EM versus light microscopy: arbor overlap is not predictive of where synapses occur. In the retina, random synapse formation on the basis of process proximity cannot explain the location of synapses found between direction-selective cells and starburst amacrine cells (Briggman et al., 2011). Similarly, the proximity of axons to dendritic spines is a poor predictor of connectivity in a densely-reconstructed ssTEM dataset spanning a volume of the mouse neocortex (Kasthuri et al., 2015).

The presence and pattern of synapses we found among DI-1, DI-102, VI-2, DE-3, and DE-4 conformed to our expectations given known physiological evidence (Ort et al., 1974). However we failed to find any synapses from DI-1, DI-102, or VI-2 onto either L cell as previous physiology predicted (Table 1) (Ort et al. 1974). The L cell is known to be electrically coupled to other excitatory motor neurons that receive direct monosynaptic inhibition from DI-1, DI-102, and VI-2 (Ort et al., 1974, Fan et al., 2005). Therefore, the synaptic input from these cells onto the L cell may be indirect while physiologically appearing otherwise (this pattern has been observed before in the leech whereby sensory cells influence the S cell via a pair of cells electrically coupled to the S cell [Muller and Scott, 1981]). This finding underscores the utility of anatomical synapse verification at the EM level: physiological connections between cells whose arbors overlap are nonetheless not necessarily monosynaptic.

Detecting electrical connections mediated by gap junctions remains an unsolved challenge in SBEM-based connectomics. In our volume, we knew certain cell pairs to be coupled, and were able to locate several places where their membranes came into prolonged contact (Figure 3). Some of these sites are almost certain to contain gap junctions, but we cannot determine how many contacts are functional versus incidental. New specimen preparation techniques (e.g., Pallotto et al., 2015) that preserve or expand the extracellular space can aid in identifying gap junctions even in SBEM. In future samples of leech neuropil, these approaches, in concert with pre or post hoc physiological verification, could lead to the description of patterns of membrane apposition associated with gap junctions in the leech.

Connectomes produce anatomical predictions of neuronal connectivity which can then be verified physiologically. In *C. elegans*, the connectome has long served as a roadmap for guiding subsequent cell ablation, imaging, and physiological experiments (Bargmann and Marder, 2013). In the larval fruit fly, connectomics predicted a neuronal circuit responsible for multisensory integration involved in rolling behavior (Ohyama et al., 2015), connections that were then verified using calcium imaging. Similarly, we demonstrated that anatomical connections can be recapitulated in physiological measurements by first discovering synapses from the S cell onto both cells 116 in our EM volume and subsequently demonstrating that spikes in the S cell produce a depolarization in cell 116 (Figure 4). This result also highlights the advantages of using an electrophysiologically accessible system in which the same cells can be identified from ganglion to ganglion and animal to animal. In principle, a complete reconstruction of a ganglion could dramatically reduce the number of pairwise recordings needed in other ganglia to confirm the existence of the identified connections, as opposed to testing every possible pair of neurons (~80,000). Importantly, in the leech ganglion these physiological experiments can involve the direct measurement of membrane potential (via intracellular electrophysiology) rather than indirect measures of activity like calcium imaging that struggle to reveal inhibitory connections.

While the leech is studied in part because of how reproducible physiological recordings are from ganglion to ganglion, anatomical features including soma position, neuronal composition (Lent et al., 1991), and fine branching patterns also vary. It is possible that there will exist some cases in which two cells are synaptically connected in some of the 21 ganglia in the nerve cord and not others, or that there are reproducible connections in different ganglia that nonetheless involve differing numbers and locations of synapses. Unfortunately, the high time and labor commitment required to produce full cell reconstructions and annotations currently limits image acquisition and analysis to a single ganglion. In other systems, measuring sample-to-sample variability from EM reconstructions has thus far been largely confined to two samples. In the earliest connectome, *C. elegans* was reconstructed from partially overlapping datasets from different animals; the connections found in the region of overlap were largely consistent from sample to sample (White et al., 1986). Similarly, in the region of overlap in two different first instar *Drosophila* larvae, 96% of connections involving two or more synapses in one animal were also found in homologous cells in the other animal (Ohyama et al., 2015), a pattern of connectivity that remained consistent in third instar larvae (Gerhard et al., 2017), though overall numbers of synapses increased proportional to the growth of the arbors. In a partial connectome of the *Platynereis* visual system, there was also a high concordance between two animals (Randel et al., 2015). Moving beyond these low N experiments will eventually require even further acceleration of imaging and analysis. In particular, automated and semi-automated reconstruction and annotation techniques currently in development (Januszewski et al., 2018; Dorkenwald et al., 2017; Staffler et al., 2017; Berning et al., 2015; Kasthuri et al., 2015; Helmstaedter 2013) could considerably decrease time costs, enabling larger sample sizes.

Our results demonstrate the utility of applying serial EM reconstruction to a system in which individual neurons can be identified from preparation to preparation. Known connections can be verified or challenged, and previously unknown connections can be discovered and subsequently tested. This connectomics approach enables the interplay between anatomical thoroughness and physiological precision that will allow future researchers to uncover previously inaccessible details regarding the circuits underpinning behavior in the leech ganglion.

## Acknowledgements

We thank Mason Mackey and Tom Deerinck for their assistance and advice for preparing and imaging the samples. This work was supported by an NIH research grant (MH43396) to WBK., an NSF grant (IOS 825741) to WBK, an NIH grant (P41GM103412) to MHE, and a grant from the Kavli Institute for Brain and Mind (to WBK and MHE). Ellisman and Bushong are carrying the leech connectomics project forward under R01 NS094403 to D. Wagenaar at Caltech and through P41GM103412.

